# Stability and flexibility in the Wisconsin Card Sorting Task emerge from goal-directed inference and planning under the free-energy principle

**DOI:** 10.64898/2026.11.17.767470

**Authors:** Daiki Goto, Hayato Idei, Tetsuya Ogata

## Abstract

In goal-directed behavior, the brain has to balance stability and flexibility. Stability means keeping the current recognition of environmental context, whereas flexibility means switching to a new one. The Wisconsin Card Sorting Task (WCST) is a standard task for studying this balance. In this task, stability is keeping the category estimate after correct feedback, and flexibility is switching it after incorrect feedback. Computational models have been proposed to elucidate the mechanism behind this balance. However, most models are given a variable for each category in advance, and every model treats correct feedback as desirable from the start. These models cover neither the category learning process nor the goal-directed process that adjusts the card selection to get correct feedback. To explain both processes under one principle, we modeled the WCST with a hierarchical recurrent neural network based on the Free Energy Principle (FEP). The model first infers the environment from observations and then plans the card selection to reach the goal. In both steps, the model minimizes prediction error based on the FEP. For inference, the error lies between the predicted signals and the received signals. For planning, it lies between the predicted feedback and the correct feedback. As a result, the latent states formed category-related clusters during learning. In the testing phase, the model kept the estimate after correct feedback and switched it after incorrect feedback. Removing the goal-directed process degraded the keeping and the switching. This implies that the two processes and this balance emerge from the FEP.

## 1. INTRODUCTION

In goal-directed behavior, a dynamic environment creates the stability–flexibility dilemma (Goschke, 2000; Dreisbach and Fröber, 2019). Stability means keep-ing the current recognition of environmental context to achieve the goal appropriately. Flexibility means switching the current recognition to a new context. The brain must balance the two. When the balance breaks, behavior becomes rigid or distractible. The purpose of this paper is to propose a mechanism behind this balance.

The Wisconsin Card Sorting Task (WCST) is a standard task for studying this dilemma (Berg, 1948; Miles et al., 2021). The participant sees one stimulus card and four target cards. Every card shows a color, a number, and a shape (Figure 1). The four target cards stay the same throughout the task, and only the stimulus card changes. The experimenter picks one of the three dimensions as the correct category. The experimenter keeps this choice secret. The participant estimates the category. The participant then selects the target card that matches the stimulus card on that category. The feedback after each choice is the only clue: correct or incorrect. After several correct selections in a row, the experimenter switches the correct category without warning. Correct feedback calls for keeping the current estimate. Incorrect feedback calls for switching it. The dilemma therefore appears as the keeping and the switching of a category estimate.

**Figure 1:**
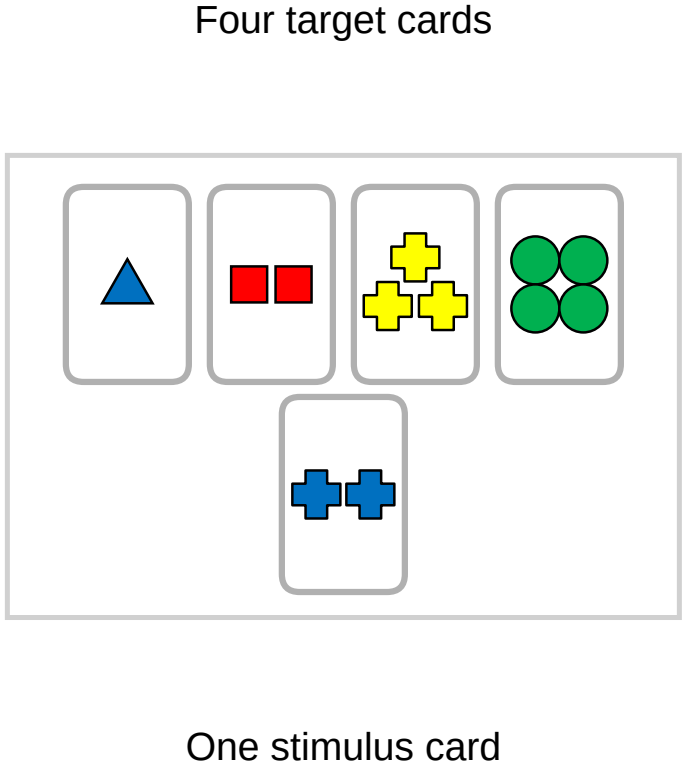
Wisconsin Card Sorting Task. One stimulus card and four target cards are presented. Every card shows a color, a number, and a shape.

In many cases, human behavior balances the two. Computational models of the WCST have been proposed to elucidate this mechanism (Levine and Prueitt, 1989; Dehaene and Changeux, 1991; Berdia and Metz, 1998; Monchi et al., 2000; Kaplan et al., 2006; Bishara et al., 2010; Caso and Cooper, 2017; Steinke et al., 2018, 2019; Caso and Cooper, 2020; Steinke et al., 2020; Steinke and Kopp, 2020; D’Alessandro et al., 2020; Granato et al., 2020; Granato and Baldassarre, 2021; Kajić and Stewart, 2021; Granato et al., 2022; Granato and Baldassarre, 2024; Liu and Wang, 2024; Granato et al., 2026). These models explain how people select the card. Also, many of them explain the card selection of patients with neurological and psychiatric disorders.

However, the WCST requires two cognitive processes that these models do not explain. The first process is to learn to distinguish the categories. The second process is to select the card as goal-directed behavior (the goal-directed process). First, most models are given a variable for each category in advance (Levine and Prueitt, 1989; Dehaene and Changeux, 1991; Berdia and Metz, 1998; Monchi et al., 2000; Kaplan et al., 2006; Bishara et al., 2010; Caso and Cooper, 2017; Steinke et al., 2018, 2019; Caso and Cooper,2020; Steinke et al., 2020; Steinke and Kopp, 2020; D’Alessandro et al., 2020; Granato et al., 2020; Granato and Baldassarre, 2021; Kajić and Stewart, 2021; Granato et al., 2022; Granato and Baldassarre, 2024; Granato et al., 2026). Therefore, these models never learn to distinguish the categories.

Second, these models treat correct feedback as desirable from the start. They are given the relation between the feedback and the category estimate in advance: the current category estimate is kept after correct feedback and moved to another category after incorrect feedback. The category estimate and the card selection are thus determined by this given relation. Some models have free parameters, but these parameters do not change the relation itself (Bishara et al., 2010; Steinke et al., 2018, 2019, 2020; D’Alessandro et al., 2020). These models do not make the estimate so that the upcoming feedback becomes correct. In other words, these models never select the card as goal-directed behavior.

Note that Liu and Wang (2024) are the only exception to the first process. Their model works on a WCST analog with two categories. It acquires the category representation through learning. However, the learning targets are always the correct category and the correct card selection. Their model therefore never selects the card as goal-directed behavior either.

The Free Energy Principle (FEP) may offer a way to explain these two processes and the balance between stability and flexibility under one principle (Friston, 2010). Under the FEP, perception, learning, and action are all explained as inferences that minimize prediction error. Prediction error is the difference between the signals the system predicts and the signals it observes. Models based on the FEP have accounted for some cognitive phenomena (Idei et al., 2022, 2025, 2026; Vijayaraghavan et al., 2025). The FEP also covers actions, including goal-directed behavior. Here the error is the difference between the predicted signals and the desired signals at future time steps. A prior belief (Friston et al., 2012) or a target signal (Matsumoto and Tani, 2020; Matsumoto et al., 2022) gives the desired signals. This formulation of action is called active inference (Friston et al., 2012). Therefore, one principle can explain learning and goal-directed behavior.

Some studies have applied the FEP to the WCST. First, Doulfoukar et al. (2026) give their model a discrete category variable and a preference for correct feedback from the start. Second, Barceló (2021) interprets the WCST based on error minimization. However, that study does not use a computational model. Third, Goto et al. (2025a) use a simplified version of the WCST. Their model sees one card and selects one of its feature values rather than one of four target cards. Therefore, no study explains the two processes based on the FEP in the standard WCST.

This paper proposes a computational model that explains the two processes based on the FEP. The model learns to distinguish the categories, and it selects the card as goal-directed behavior. Also, the balance between stability and flexibility emerges from this goal-directed process. We model the WCST with a hierarchical recurrent neural network (RNN) based on the FEP, and we run simulation experiments with the model.

To model the WCST, we have to cover the cognitive requirements that the task places on the participant. Before the task, the participant learns the relation among the cards, the card selection, and the feedback. During the task, the participant perceives the cards. The participant produces a card selection that aims at correct feedback. The participant receives the feedback and revises the category estimate. The model works in two phases. During learning, the model predicts the cards, the card selection, and the feedback. The model minimizes the error between these predictions and the observed signals. The model thus learns the relation among the three signals. Through this learning, the model comes to distinguish the categories (the category learning process). In this phase, the model does not treat correct feedback as desirable. During the testing phase, the model first receives the cards as observed signals. Second, the model treats correct feedback as the target signal at future time steps. The model adjusts its internal state to minimize the error between the target and the predicted feedback at future time steps, and the card selection follows from this state. This is the goal-directed process. Third, the model receives the feedback on the card selection. The model adjusts the internal state to minimize the error between the predicted feedback and the observed feedback.

The learning experiments show that the model distinguishes the three categories (Section 3.1). The test experiments show that the model keeps the estimate after correct feedback and switches it after incorrect feedback (Section 3.2). Removing the goal-directed process degrades both the keeping and the switching (Section 3.3). Therefore, the balance between stability and flexibility emerges from prediction error minimization based on the FEP in the goal-directed process.

In summary, in a hierarchical RNN given neither a variable for each category nor a preference for correct feedback, the two cognitive processes and the balance between stability and flexibility emerge from prediction error minimization based on the FEP. This implies that cognitive processes and this balance in the WCST emerge from the FEP.

## 2. EXPERIMENT & PROPOSED MODEL

In this study, we conducted simulation experiments of the WCST using an RNN model based on the FEP. The experiments consisted of a learning phase, in which the model learned from prepared sequences, and a testing phase, in which the trained model selected cards by itself. A single trial in the experiment consisted of four phases (Figure 2):

- Phase 1 (Card recognition): Cards were presented to the model
- Phase 2 (Card selection): The model selected one of the four target cards
- Phase 3 (Feedback): A feedback signal was presented to the model
- Phase 4 (Reset): The model returned to the initial state

**Figure 2:**
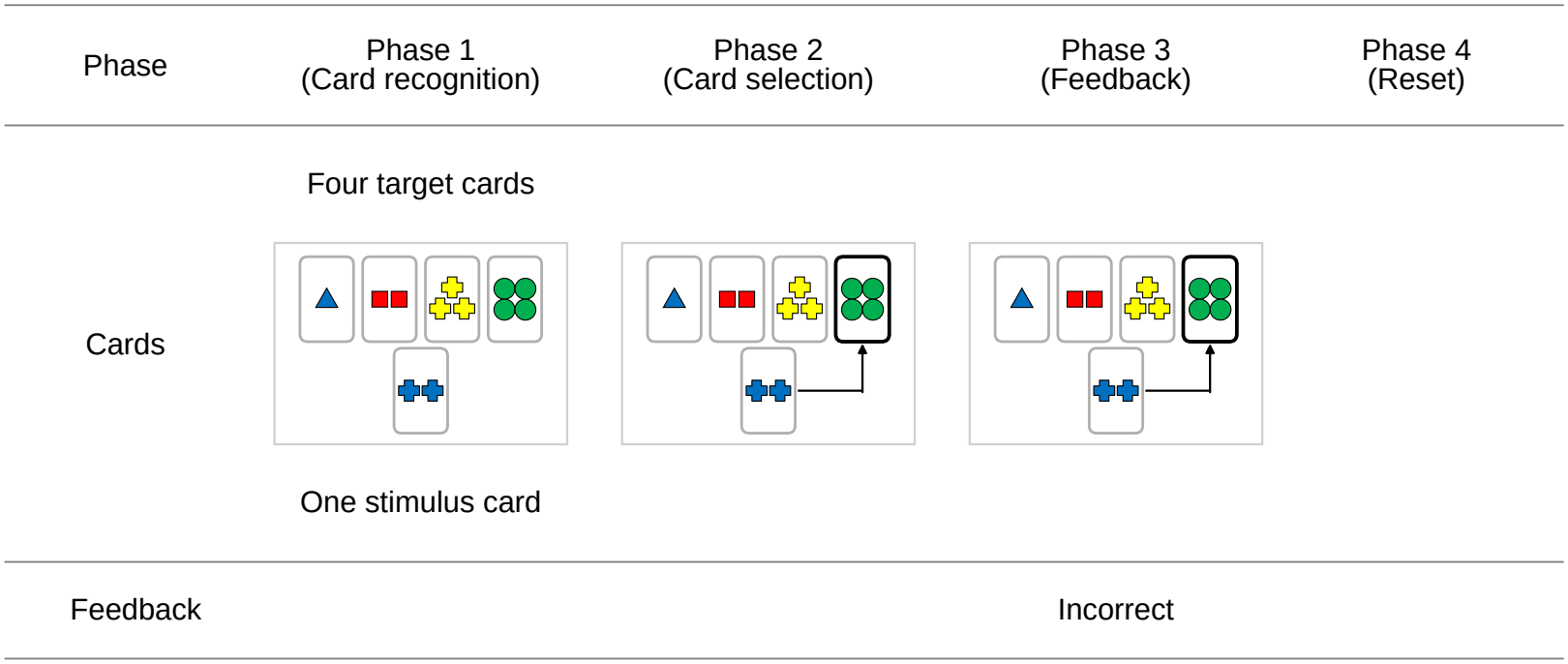
Overview of a single trial in the WCST simulation experiment. Phase 1 presented five cards (one stimulus card and four target cards). In Phase 2, the model selected a target card. Phase 3 provided feedback on the card selection. Phase 4 returned the model to the initial state.

Each phase consisted of 16 time steps, and a single trial consisted of 64 time steps in total. Figure 3 shows how the signals changed within a trial: the card signals turned on in Phase 1, the card selection signal in Phase 2, and the feedback signal in Phase 3, and all signals returned to zero in Phase 4.

**Figure 3:**
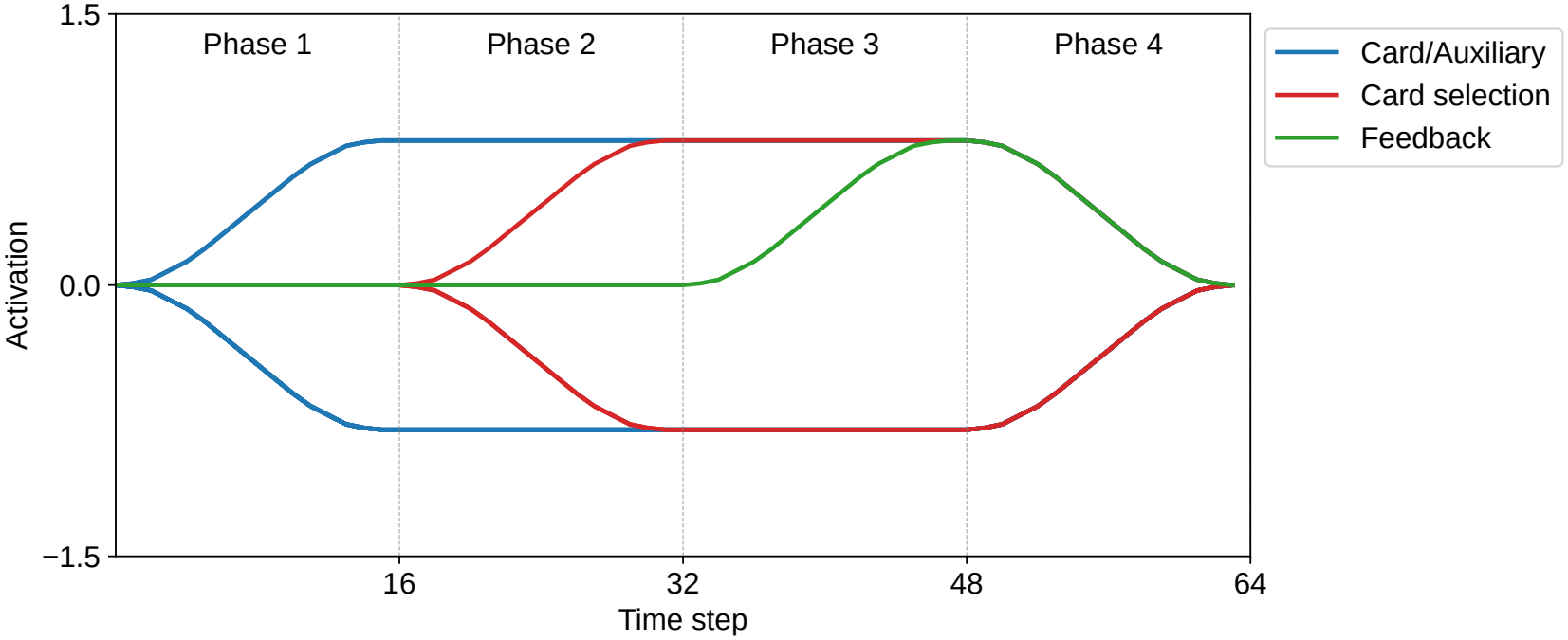
Temporal activation schedule within a single trial. Each trial consisted of 64 time steps divided into four phases (16 steps per phase). The figure shows a trial with correct feedback. Card information turned on during Phase 1, the card selection signal during Phase 2, and the feedback signal during Phase 3. Each signal stayed on until Phase 4, where all signals returned to zero. The card signals and the card selection signal were one-hot vectors, so the figure plots one example of a signal that took 0.8 and one example of a signal that took *−*0.8. The feedback signal was one-dimensional, and it took 0.8 in a trial with correct feedback. Values changed smoothly at the start of each phase and in Phase 4. Auxiliary information is described in Section 2.2.

Within a single trial, the proposed model predicted the following multi-dimensional time-series data as sensory signals:

- Five cards (one stimulus card and four target cards), each bearing features related to color, number, and shape
- Card selection
- Correct/incorrect feedback

The four target cards were the same in the learning phase and the testing phase.

### 2.1. Proposed Model

We modeled the WCST with a two-layer Predictive-coding-Inspired Variational Recurrent Neural Network (PV-RNN; Ahmadi and Tani, 2019; Idei et al., 2022) (Figure 4). The PV-RNN is an RNN based on the FEP that infers the hidden causes of the sensory signals. The network has deterministic states *d* and latent states *z*. The latent states hold beliefs about the hidden causes as Gaussian distributions. Each latent state has a prior *z*^*p*^ and a posterior *z*^*q*^. The prior is the estimate before the network observes the signals. The posterior is the estimate after the observation. The network infers the posterior by minimizing variational free energy. This free energy is the sum of the prediction error (PE) and the Kullback–Leibler divergence (KLD) between the posterior and the prior. Using these deterministic states and latent states, the network generates predictions about the sensory signals at the next time step.

**Figure 4:**
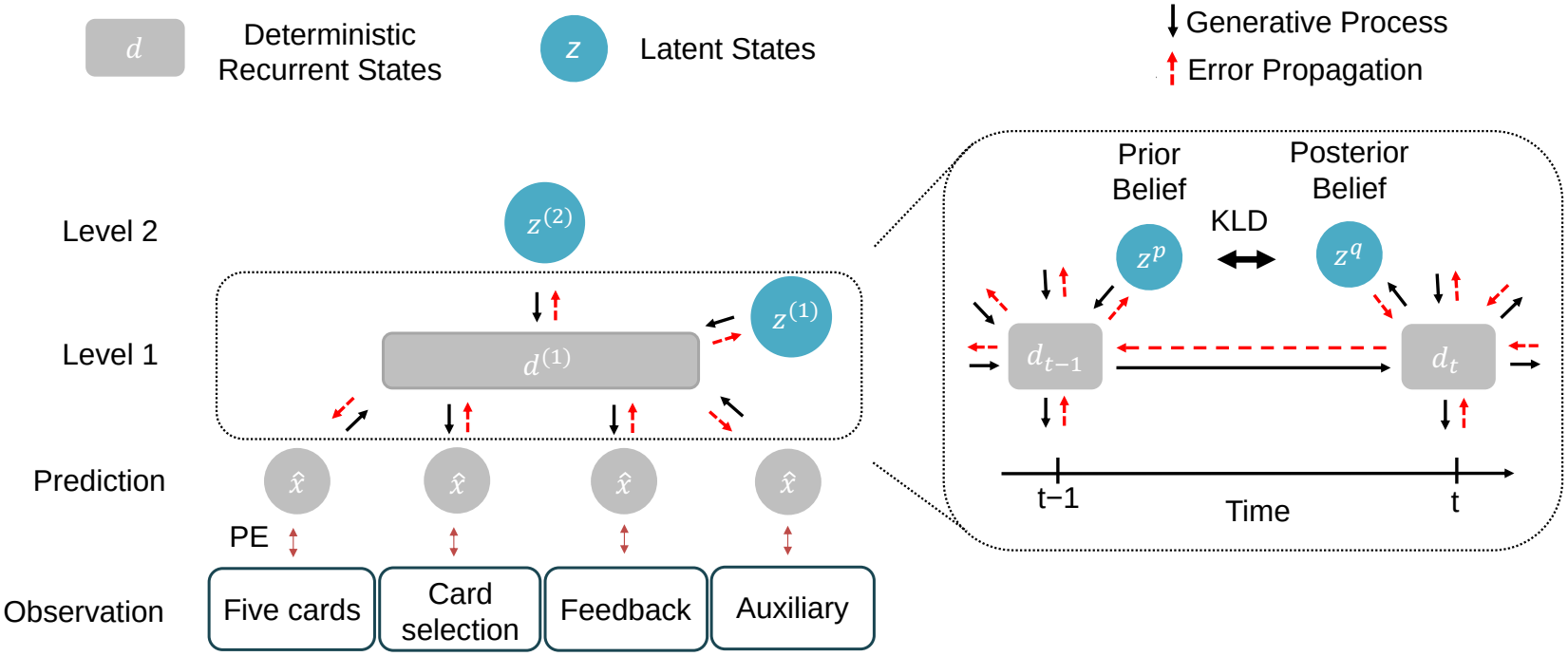
Structure of the two-layer Predictive-coding-Inspired Variational Recurrent Neural Network (PV-RNN). The network has deterministic states *d* and latent states *z*. The latent states represent beliefs as Gaussian distributions with a prior *z*^*p*^ and a posterior *z*^*q*^. The network generates predictions of the sensory signals from these states. PE denotes the prediction error between the predicted and the observed sensory signals. KLD denotes the Kullback–Leibler divergence between the posterior and the prior. Auxiliary information is described in Section 2.2. The superscripts (1) and (2) denote the first and second layers.

#### 2.1.1. Prediction Generation

Predictions are generated in a top-down manner through the network hierarchy. At time step *t ≥* 1, the internal state 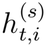 and output 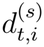 of the *i*-th deterministic state in the first layer in the *s*-th sequence are computed as follows:

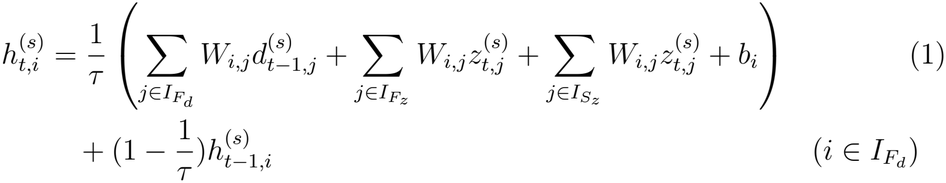

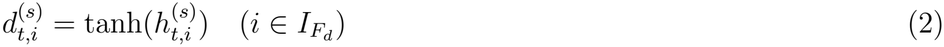

Here, 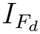 denotes the set of indices of deterministic states in the first layer, and 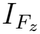 and 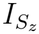 denote the sets of indices of latent states in the first and second layers, respectively. *W*_*i,j*_ represents the synaptic weight from the *j*-th state to the *i*-th state, and *b*_*i*_ is the bias of the *i*-th state. *z*_*t,j*_ represents the output of the posterior distribution of the *j*-th state at time step *t*. The time constant *τ* controls the temporal scale of each state’s activity; a small *τ* results in faster activity changes, and a large *τ* results in slower changes. The initial internal state 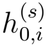 of deterministic states is zero.

Latent states are represented as independent multivariate Gaussian distributions in each layer. The prior distribution 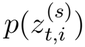 is computed from the deterministic states at the previous time step in the same layer using the mean *µ* and standard deviation *σ* as follows:

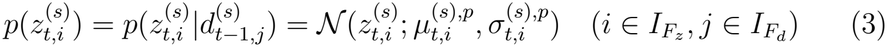

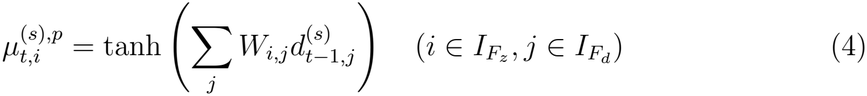

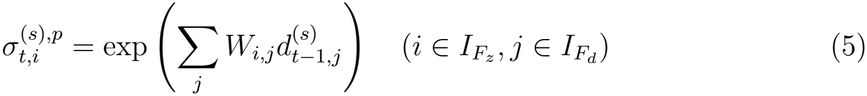

The latent states in the second layer maintain constant posterior distribution values within each sequence, with the goal of assigning specific posterior distribution values to each learning sequence. Because of this constant posterior, their prior is defined only at the initial time step (*t* = 1) as *N* (0, 1).

The posterior distribution *z*^*q*^ is computed as follows:

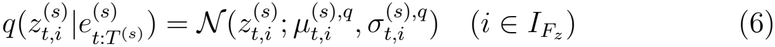

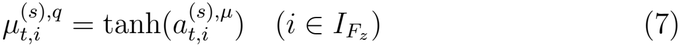

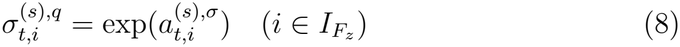

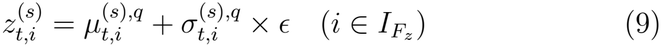

Here, *T* ^(*s*)^ denotes the length of data in sequence *s*. Additionally, *ϵ* is sampled from *N* (0, 1). Also, 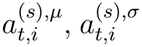 are adaptive variables that represent the posterior distribution of the corresponding state; they are updated at each time step for each sequence during both the learning and testing phases.

Equations (7) and (8) imply that the adaptive variables implicitly encode both the mean and precision of the posterior distribution, and that optimizing the adaptive variables with respect to variational free energy implicitly optimizes the mean and precision of the posterior distribution. The adaptive variables 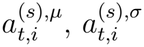 are determined by the prediction error signal 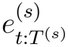 ropagated by the backpropagation through time (BPTT) algorithm.

The adaptive variables 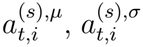 are initialized by the initial internal state of latent states representing the prior distribution (the value before passing through the activation function tanh or exp).

Finally, the predictive signals for each type of sensory information are generated from the deterministic states in the first layer. Let *I*_*O*_ denote the index set of the predictive signals.

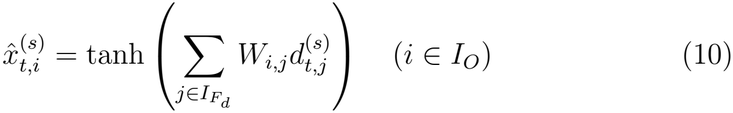

Here, 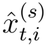 denotes the *i*-th predictive signal at time step *t*. Predictive signals 1–85 corresponded to the five cards. Each card was represented as a 17-dimensional vector: four dimensions each for color, number, and shape, and five dimensions for the card index (1–5). Signals 86–89 corresponded to the card selection, represented by a 4-dimensional one-hot vector specifying which of the four target cards was selected. The dimension with the maximum value at the card selection phase gave the selected card. Each one-hot vector took a value of 0.8 for the active dimension and *−*0.8 for the other dimensions. Signal 90 corresponded to correct/incorrect feedback, with a value of 0.8 for correct card selection and *−*0.8 for incorrect card selection.

### 2.2. Learning

The goal of the learning phase was to predict time-series data of the learning sequences. Each learning sequence consisted of eight trials, that is, 512 time steps. Category switches were not included within the learning sequences. The learning data consisted of 54 sequences in total: nine sequences for each of the three categories with correct card selection, and nine sequences for each category with incorrect card selection.

Each network updated synaptic weights and adaptive variables to minimize the variational free energy *F*_learning_ across all time steps and learning sequences. We trained 10 networks with different random initializations. For notational simplicity, we omit the state index *i* and the sequence index *s* in the Kullback–Leibler divergence terms below.

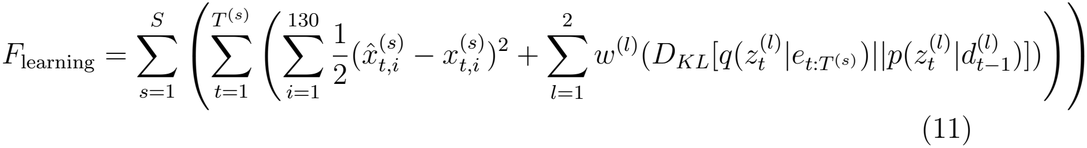

The first term is the prediction error term and the second term is the Kullback–Leibler divergence term. Here, *t* denotes the time step, *l* is the layer index, *s* is the sequence index, *S* is the number of sequences in the data, and 130 is the dimensionality of the sensory signals. Furthermore, *d* denotes the output of deterministic states, *z* denotes the value of latent states, and *p* and *q* denote the prior and posterior distributions, respectively. Additionally, *e* denotes the prediction errors from time step *t* to the final time step *T* ^(*s*)^ of sequence *s*. The parameter *w*, called the meta-prior, weights the first and second terms.

To select a card, the model has to group the features from the five cards by category and pay attention only to the features it needs. This ability is called dimensional selective attention, and it is acquired through development in humans (Hanania and Smith, 2010). The learning of words for color and shape has been reported to correlate with this development (Lowery et al., 2022). Learning such a word means extracting a single dimension from the object. Motivated by this observation, in the learning phase we extracted the card features along particular category dimensions and gave them as sensory signals in addition to the five cards. These signals formed the auxiliary information, and they were intended to support the acquisition of dimensional selective attention. The auxiliary information consisted of two components:

- Card features corresponding to the correct category dimension
- Card features corresponding to the attended category dimension

We refer to the category that the card selection follows as the attended category because it stands for the dimension that the model attends to when it selects a card. Consider an incorrect card selection sequence. The correct category was shape, and the card selection followed color (the attended category). In this case, we additionally provided the shape information of each card (4-dimensional one-hot vector *×* five cards) and the color information of each card (4-dimensional one-hot vector *×* five cards) as sensory signals. In a correct card selection sequence, the two components carried the same card features, because the correct category and the attended category agreed. The auxiliary information thus consisted of card features along the relevant dimensions, and it did not give the categories as discrete labels. Accordingly, the sensory signals comprised 130 dimensions in total, where signals 91–110 corresponded to auxiliary information about the correct category and signals 111–130 corresponded to auxiliary information about the attended category. We provided the auxiliary information only in the learning phase, and we removed it in the testing phase. In the testing phase, the model received only the 90 dimensions that corresponded to the five cards, its own card selection, and the feedback.

#### 2.2.1. Error Scheduling

Learning the task involves several kinds of information processing: perceiving the cards, extracting the features along the correct category, extracting the features along the attended category, and relating the card selection to the feedback. To let the model acquire them step by step, we used error scheduling. We added a new group of dimensions to the error signal every 5,000 epochs in the order listed (Table 1). At the end of learning, all dimensions contributed to the prediction error.

**Table 1:** Error scheduling during learning. Error signals were progressively added at 5,000-epoch intervals.

| **Training Phase** | **Information Added** |
|---|---|
| 0 epochs | (1) Card features (five cards) |
| 5,000 epochs | (2) Auxiliary information of the correct category |
| 10,000 epochs | (3) Auxiliary information of the attended category |
| 15,000 epochs | (4) Card selection and feedback signals |

### 2.3. Test

In the testing phase, we used the trained networks to generate card selection for novel test sequences that were not provided as learning data. Unlike the learning data, the correct category switched within the test sequences. At this stage, synaptic weights were fixed to their learned values and only adaptive variables were updated.

Card selection in the WCST was implemented based on free energy minimization utilizing past and future windows (Figure 5). At each current time step *t*_*c*_, the model updated the adaptive variables within the range from *t*_*c*_ *−win*_*p*_+1 to *t*_*c*_+*win*_*f*_ to minimize the total free energy *F*_test_ = *V*_past_+*G*_future_, where *win*_*p*_ and *win*_*f*_ denote the lengths of the past and future windows, respectively. The windows slid forward as *t*_*c*_ progressed.

**Figure 5:**
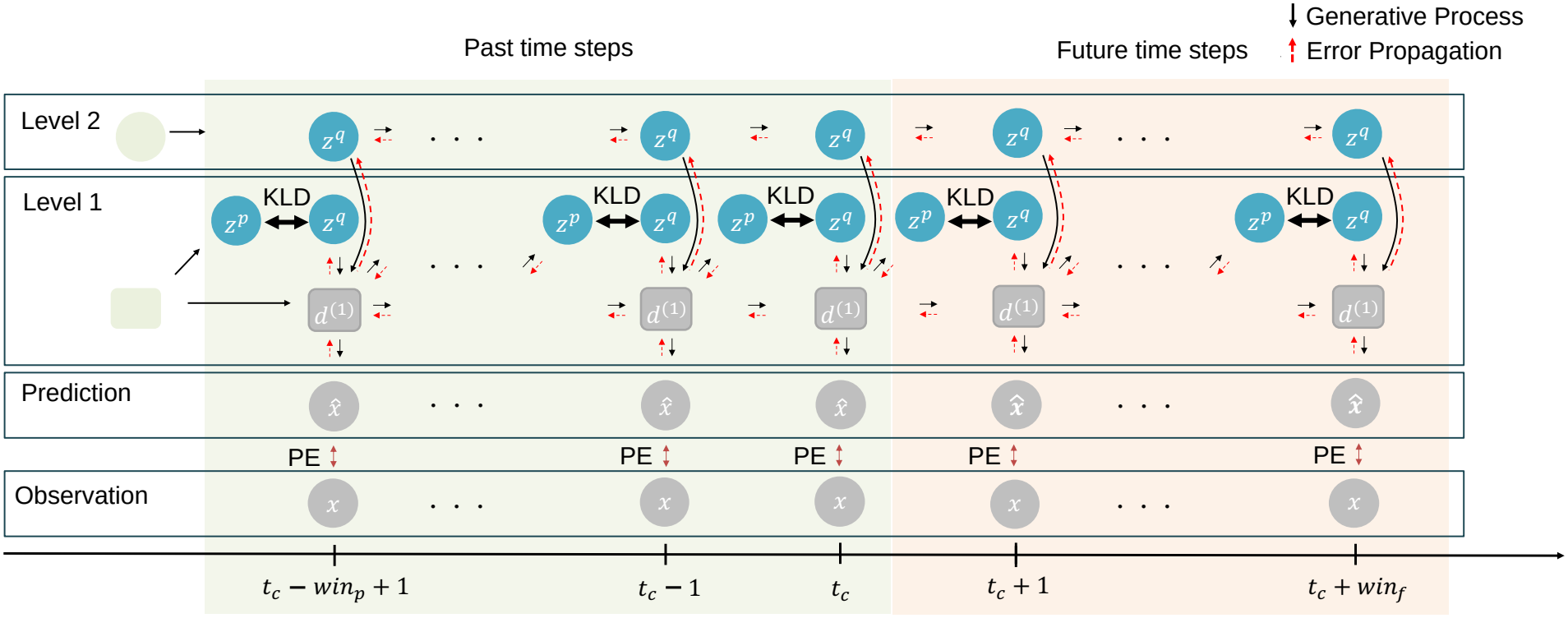
Free energy minimization with past and future windows. At the current time step *t*_*c*_, the model updated adaptive variables within the range from *t*_*c*_ *−win*_*p*_ + 1 to *t*_*c*_ + *win*_*f*_, where *win*_*p*_ and *win*_*f*_ denote the lengths of the past and future windows, respectively.

In the past window (*t*_*c*_ *−win*_*p*_ + 1 to *t*_*c*_), the prediction error 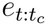 was computed as the error between the observed sensory signals and the generated predictive signals for the five cards, the model’s own card selection, and the external feedback (Figure 6a). The variational free energy *V*_past_ was computed as follows:

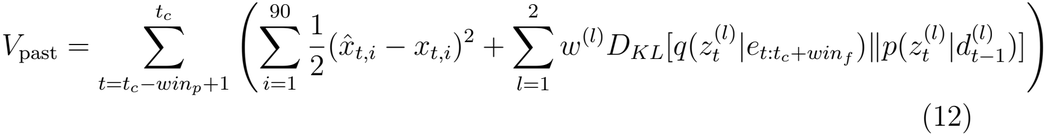

**Figure 6:**
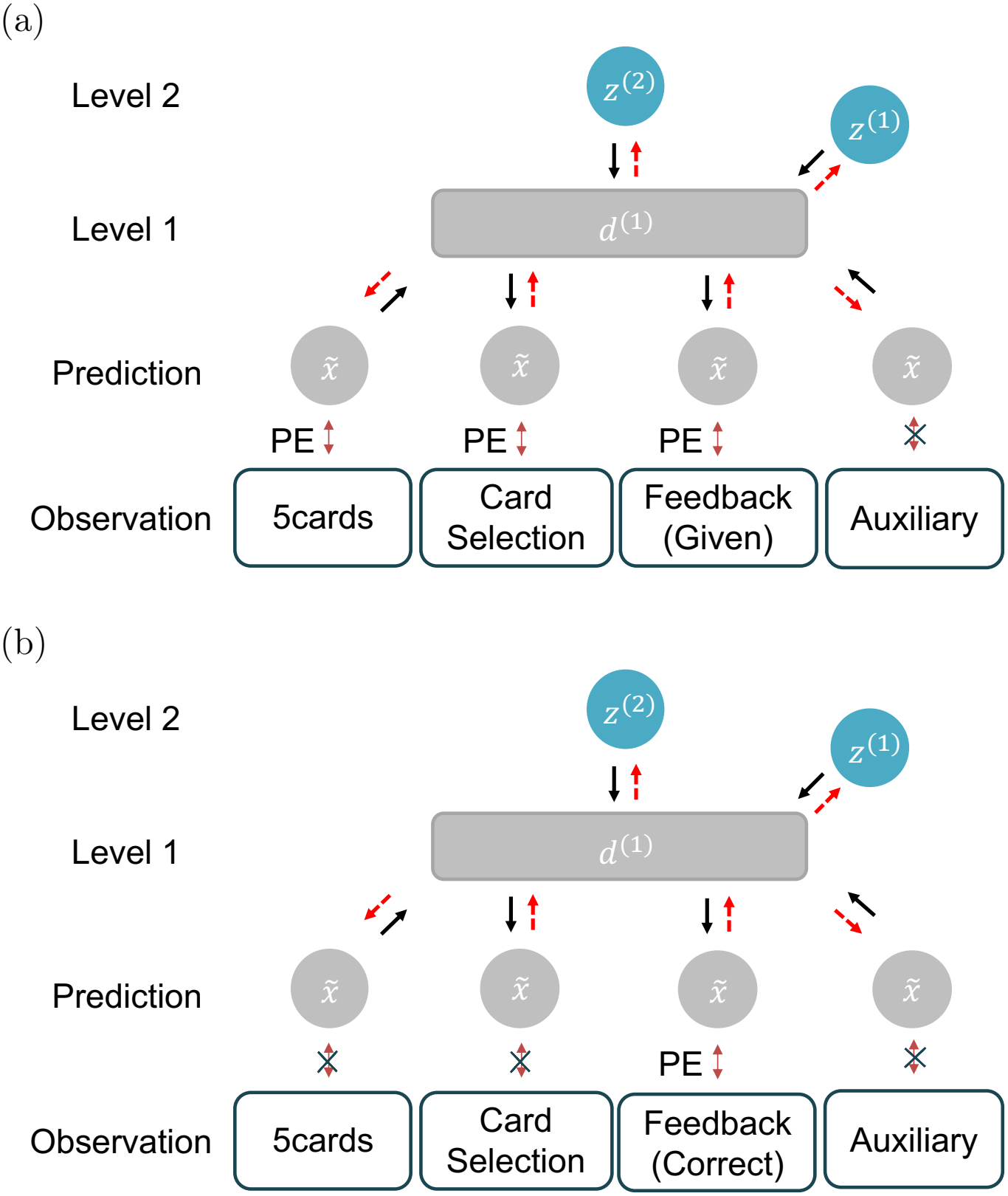
Free energy minimization at past and future time steps. (a) The model computed the variational free energy *V*_past_ over past time steps. It minimized the error between the observed sensory signals and the predictive signals. (b) The model computed the future free energy *G*_future_ over future time steps. It minimized the error between correct feedback, which was the target signal, and the predicted feedback.

Here, the first term is the prediction error term and the second term is the Kullback–Leibler divergence term. The prediction error term sums over the 90 dimensions (*i* = 1, …, 90) corresponding to the five cards, the card selection, and the feedback.

In the future window (from *t*_*c*_ + 1 to *t*_*c*_ + *win*_*f*_), we gave the model correct feedback as the target signal. This target served as the goal for the feedback. We gave no target for the other dimensions. The prediction error 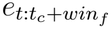 was computed as the error between the target signal and the predicted feedback (Figure 6b). The future free energy *G*_future_ was computed as follows:

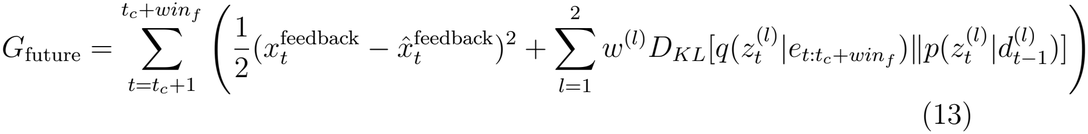

By minimizing *G*_future_, the model adjusted its internal state at the current time step so that the predicted feedback approached correct feedback, and the card selection followed from this state. This minimization of *G*_future_ over the future window is the goal-directed process of the model.

### 2.4. Parameter Settings

The first layer had 200 deterministic states. The first and second layers had five and two latent states, respectively. We set the time constant *τ* in the first layer to 2.0 for half of the 200 deterministic states and to 4.0 for the other half, and the meta-prior to *w* = 0.005 in all layers.

In the learning phase, we trained the networks for 20,000 epochs. We used Rectified Adam with *α* = 0.001 (the learning rate), *β*_1_ = 0.9, and *β*_2_ = 0.999, and applied a weight decay of 0.0001 to the synaptic weights.

In the testing phase, we selected the past and future window lengths from {20, 30, 50} time steps, the number of posterior updates at each time step from {20, 30, 50}, and *α* for Rectified Adam from {1.0, 5.0, 10.0}. We ran all combinations of these values and adopted the combination with the highest correct selection rate for each test sequence. When several combinations gave the same highest rate, we adopted the one tested first.

## 3. RESULTS

### 3.1. Learning of the category distinction

The first process is the category learning process. To evaluate this process, we examined the latent states in the second layer at the end of the learning phase.

Figure 7 plots these latent states for a representative network. We trained 10 networks with different random initializations and selected one of them for the plot. The sequences separated according to the correct category, that is, color, number, or shape. The network received no category label during learning, although we gave the network auxiliary signals that carried the card features of the correct category and of the attended category (Section 2.2).

**Figure 7:**
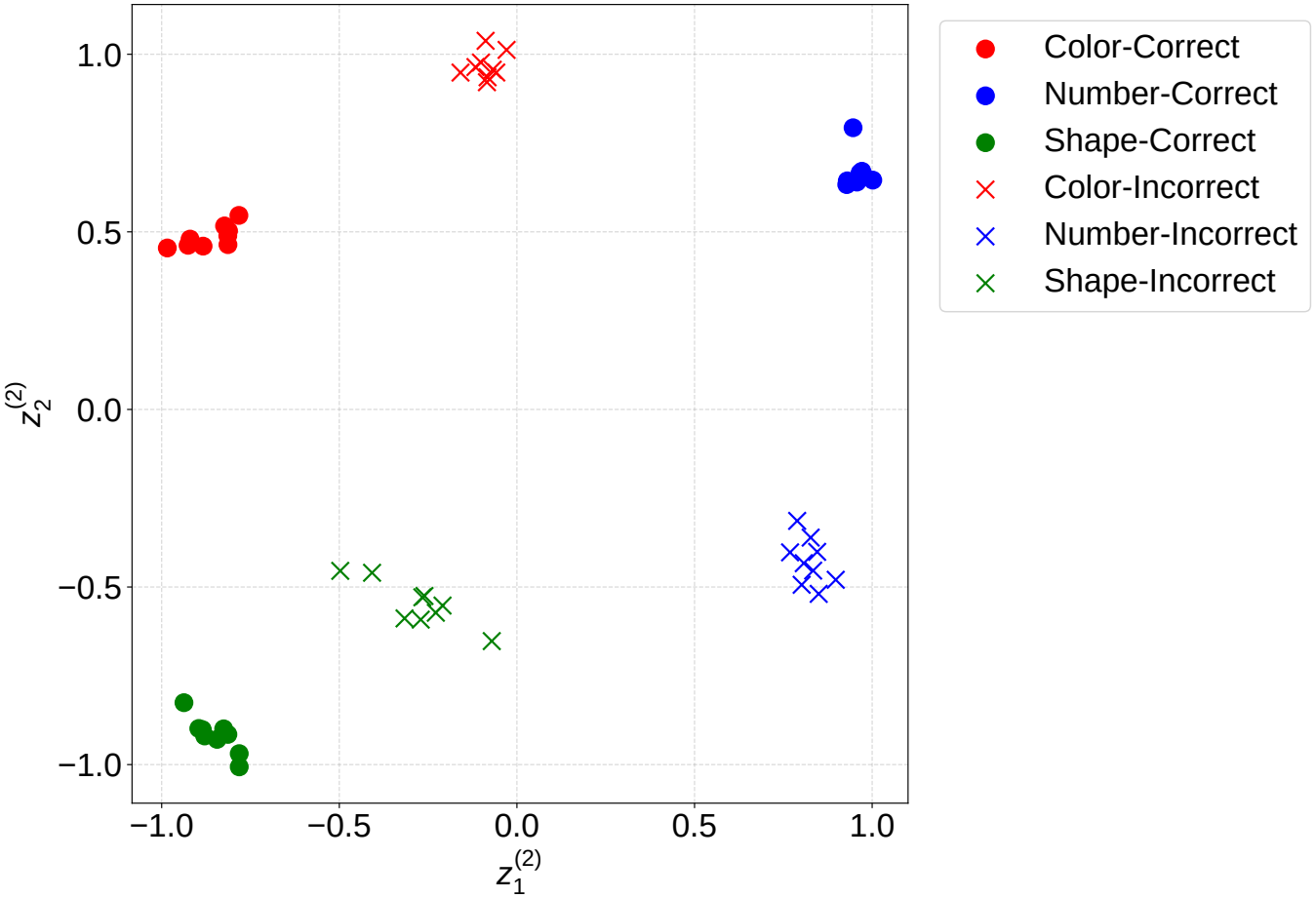
Latent space mapping of the second layer in a representative network. The latent states in the second layer 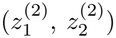 are mapped at the end of the learning phase. The network shown is one of 10 networks trained with different random initializations. Filled circles (•) represent sequences given correct feedback signals, while crosses (*×*) represent sequences given incorrect feedback signals. Red, blue, and green plots correspond to sequences where the correct category was color, number, and shape, respectively.

We then quantified the separation. We applied k-means clustering to the second-layer latent states of each of the 10 networks. We computed the within-cluster sum of squares (WCSS) for k = 2 to 10. The WCSS is the sum of squared distances between each data point and its cluster centroid. We selected the optimal k with the elbow method. The method detects the inflection point of the WCSS curve from its second-order difference. Figure 8 marks the optimal k for each network with a red star.

**Figure 8:**
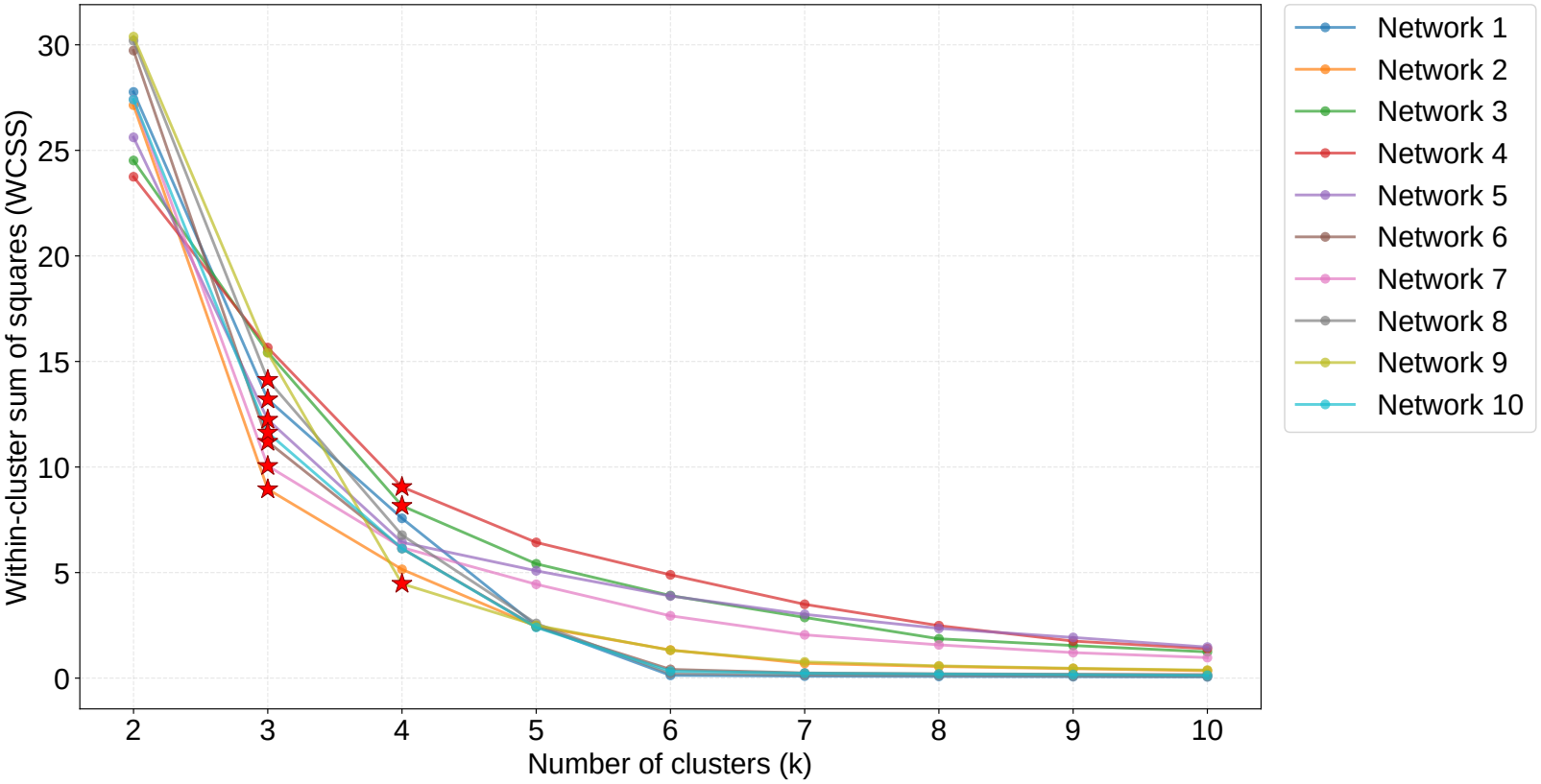
Elbow curves for k-means clustering of the second-layer latent states, computed separately for each of the 10 trained networks. For each network, the within-cluster sum of squares (WCSS) was computed for k = 2 to 10, and the optimal number of clusters was identified using the elbow method (red stars). The elbow method selected *k* = 3 for seven networks and *k* = 4 for three networks.

The elbow method selected *k* = 3 for seven of the 10 networks and *k* = 4 for the other three. In these seven networks, the second-layer latent states therefore formed three clusters, which may correspond to the three categories (color, number, and shape; Figure 7). Learning to distinguish the categories may therefore have emerged from prediction error minimization based on the FEP in the learning phase.

### 3.2. Keeping and switching of the category estimate under feedback

The second process is to select the card as goal-directed behavior. In this process, the model selects the card from its category estimate, and it has to keep the estimate after correct feedback and switch it after incorrect feedback. To evaluate this process, we ran the trained networks on a test sequence with category switches. We set the hyperparameters as follows: a future window length of 50, a past window length of 20, 20 posterior updates per time step, and *α* = 5.0 for Rectified Adam.

Figure 9 shows the card selection and the latent state dynamics of a representative network for one sequence. The sequence consisted of 20 trials. The experimenter switched the correct category after three consecutive correct card selections, and the sequence contained three switches (Top row).

**Figure 9:**
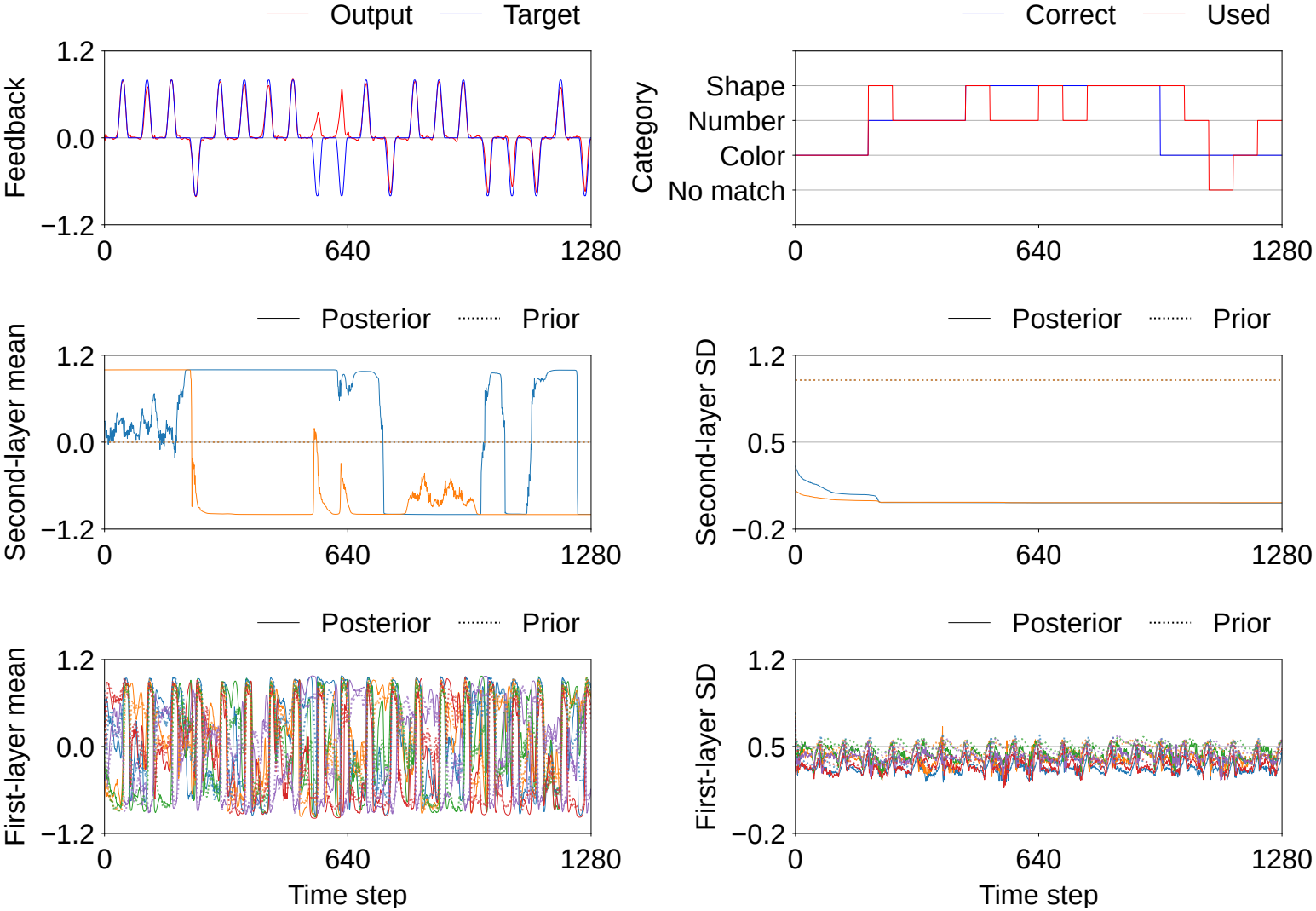
Network output and latent state dynamics for a representative test sequence showing category switching (1,280 time steps; 20 trials with three category switches).*Top row:* (Left) Feedback signal: blue = target, red = network’s output (predicted feed-back). (Right) Category estimates: blue = correct category; red = category used by the network. No match denotes a card selection that matches the stimulus card on none of the three categories. *Middle row:* Second-layer latent states (2-dimensional): (Left) Mean of posterior and prior. (Right) Standard deviation (SD) of posterior and prior. *Bottom row:* First-layer latent states (5-dimensional): (Left) Mean of posterior and prior. (Right) Standard deviation of posterior and prior. In the Middle and Bottom rows, each line shows one latent state; solid lines show the posterior and dotted lines show the prior.

The posterior mean of the second-layer latent states moved in response to the feedback signals, and the network reached the correct category after each switch (Middle row). The category used in the card selection followed the estimate, and the network recovered the correct card selection within a few trials after each switch (Top row). In contrast, the first-layer latent states changed within each trial rather than in response to the feedback signals (Bottom row). They appeared to process the sensory signals sequentially. In summary, the model selected the card from its category estimate, kept the estimate after correct feedback, and switched it after incorrect feedback.

### 3.3. Contribution of the goal-directed process to the keeping and the switching

We quantitatively investigated whether the goal-directed process produced the keeping and the switching. We compared the proposed model with a model without the goal-directed process. The latter minimized the error at past time steps only.

To compare the two conditions under the same settings, we fixed the number of trials, the number of category switches, and the timing of the switch in each sequence. Each sequence consisted of 20 trials with one category switch, and the experimenter switched the category after 10 trials. The switching patterns covered all ordered pairs of the three categories. This gave six patterns. We used three sequences per pattern, which gave 18 sequences in total.

At the start of each sequence, we set the second-layer latent state to its value at the end of learning for a learning sequence with correct card selection for the first correct category.

We evaluated the model with three metrics derived from its card selection. The correct selection rate is the proportion of trials in which the selected card matched the stimulus card on the correct category. The perseverative error rate is the proportion of trials after incorrect feedback in which the model selected the card by the same category as in the previous trial. It reflects a lack of flexibility. The non-perseverative error rate is the proportion of trials after correct feedback in which the model selected the card by a different category from that in the previous trial. It reflects a lack of stability.

Figure 10 compares the three metrics between the networks with and without the goal-directed process at the behavioral level. The networks with the goal-directed process gave a higher correct selection rate and lower error rates.

**Figure 10:**
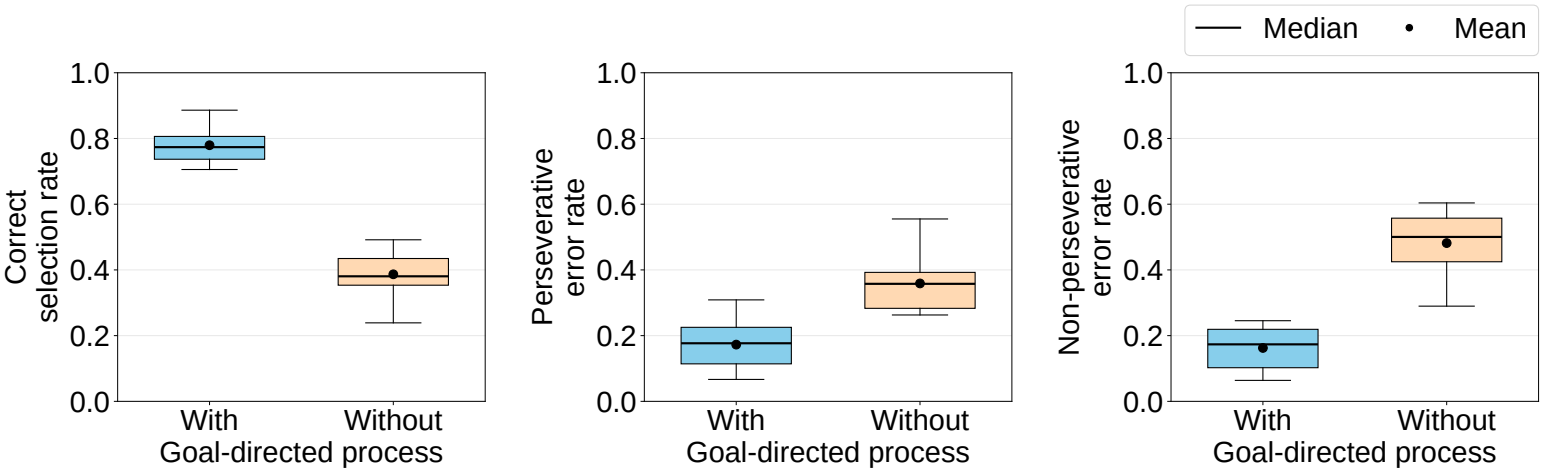
Comparison of evaluation metrics (correct selection rate, perseverative error rate, and non-perseverative error rate) at the behavioral level, between the networks with and without the goal-directed process. Each box summarizes 10 networks.

Also, we computed the same three metrics based on the second-layer latent states. These metrics show whether the model estimated the category correctly, independently of its card selection. At each trial, we determined the category that the latent state pointed to. For this determination, we used the value of the second-layer latent state of each learning sequence at the end of learning. The latent space separated not only by category but also by whether the card selection was correct, so we restricted these values to the 27 correct card selection sequences, nine for each category. At each trial of the test sequences, we found the value closest to the latent state among these 27 values, and we took the category of the corresponding learning sequence as the category that the network estimated. The latent-state correct selection rate is the proportion of trials in which the estimated category matched the correct category. The latent-state perseverative error rate is the proportion of trials after incorrect feedback in which the estimated category was the same as in the previous trial. The latent-state non-perseverative error rate is the proportion of trials after correct feedback in which the estimated category differed from that in the previous trial.

Figure 11 compares the three metrics between the networks with and without the goal-directed process at the latent state level. The networks with the goal-directed process gave a higher correct selection rate and lower error rates at both the behavioral and latent state levels.

**Figure 11:**
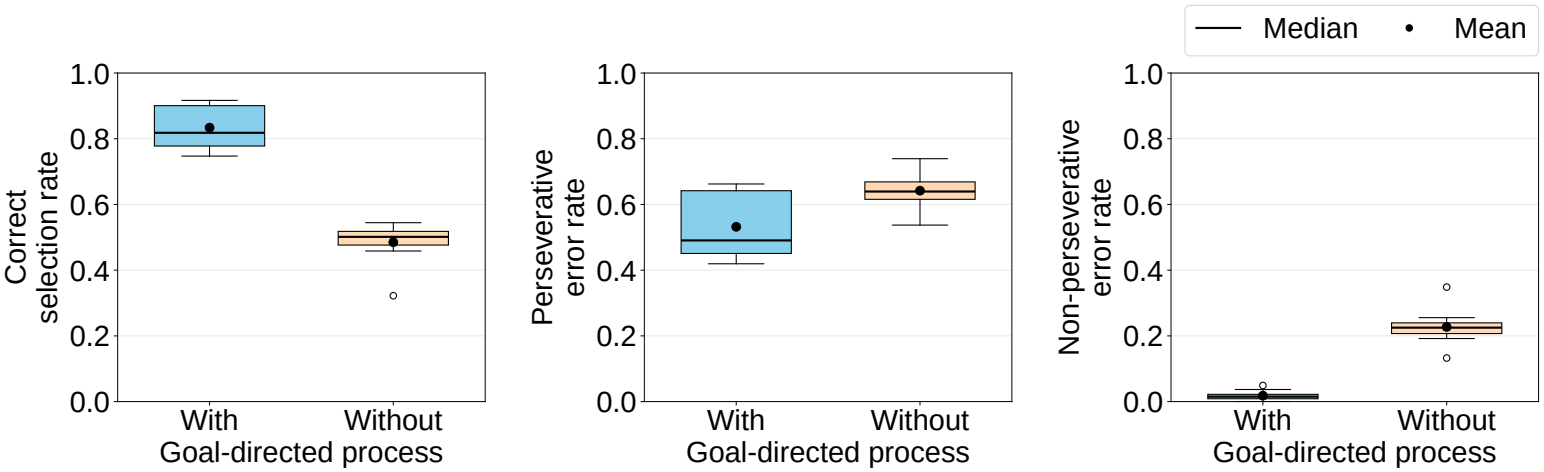
Comparison of evaluation metrics (correct selection rate, perseverative error rate, and non-perseverative error rate) at the latent state level, between the networks with and without the goal-directed process. Each box summarizes 10 networks.

Table 2 reports paired t-tests over the 10 networks with different random initializations, in which each metric was averaged over the 18 sequences for each network. For evaluation, we explored all hyperparameter combinations for the networks with the goal-directed process and adopted the combination with the highest correct selection rate for each sequence. We applied the same hyperparameter combination to the networks without the goal-directed process. All six differences were significant: five at *p* < .001 and the latent-state perseverative error rate at *p* < .005. These results did not change after Holm correction for the six comparisons.

**Table 2:** Paired t-tests comparing the networks with and without the goal-directed process. Positive *t* values indicate higher values for the networks with the goal-directed process. Each metric was computed at both the behavioral and latent state levels, across 10 networks with different random initializations evaluated on 18 sequences.

| **Metric** | *t*(9) | ***p*-value** | **Sig.** |
|---|---|---|---|
| *Behavioral Metrics* |  |  |  |
| Correct selection rate | 14.8 | 1.3 × 10<sup>−7</sup> | *** |
| Perseverative error rate | −6.0 | 2.1 × 10<sup>−4</sup> | *** |
| Non-perseverative error rate | −9.0 | 8.9 × 10<sup>−6</sup> | *** |
| *Latent State Metrics* |  |  |  |
| Correct selection rate | 13.7 | 2.5 × 10<sup>−7</sup> | *** |
| Perseverative error rate | −3.8 | 4.2 × 10<sup>−3</sup> | ** |
| Non-perseverative error rate | −12.0 | 7.7 × 10<sup>−7</sup> | *** |

Note: *p*-values are uncorrected. \*\* denotes *p* < .005, \*\*\* denotes *p* < .001. All results remained significant at the same levels after Holm correction for the six comparisons.

The balance between stability and flexibility therefore emerged from prediction error minimization based on the FEP in the goal-directed process.

#### 3.4. Contribution of the error scheduling to the keeping and the switching

We investigated whether the error scheduling contributed to the keeping and the switching. We ran the comparison with networks trained without the error scheduling. In the networks without the error scheduling, all error signals were provided from the outset of learning. The setup and the measures were the same as in Section 3.3. We also used the same hyperparameter combinations and the same initial weights as in the networks with the error scheduling.

Figure 12 compares the six metrics between the networks with and without the error scheduling. The networks with the error scheduling gave a higher correct selection rate and lower error rates at both levels. Table 3 reports paired t-tests over the 10 networks in the same way. All six differences were significant (*p* < .001). These results did not change after Holm correction for the six comparisons.

**Table 3:** Paired t-tests comparing the networks with and without the error scheduling. Positive *t* values indicate higher values for the networks with the error scheduling. Each metric was computed at both the behavioral and latent state levels, across 10 networks with different random initializations evaluated on 18 sequences.

| **Metric** | *t*(9) | ***p*-value** | **Sig.** |
|---|---|---|---|
| *Behavioral Metrics* |  |  |  |
| Correct selection rate | 15.4 | 8.9 × 10<sup>−8</sup> | *** |
| Perseverative error rate | −5.8 | 2.6 × 10<sup>−4</sup> | *** |
| Non-perseverative error rate | −6.7 | 9.3 × 10<sup>−5</sup> | *** |
| *Latent State Metrics* |  |  |  |
| Correct selection rate | 15.8 | 7.2 × 10<sup>−8</sup> | *** |
| Perseverative error rate | −8.0 | 2.3 × 10<sup>−5</sup> | *** |
| Non-perseverative error rate | −6.5 | 1.2 × 10<sup>−4</sup> | *** |

Note: *p*-values are uncorrected. \*\*\* denotes *p* < .001. All results remained significant at the same levels after Holm correction for the six comparisons.

**Figure 12:**
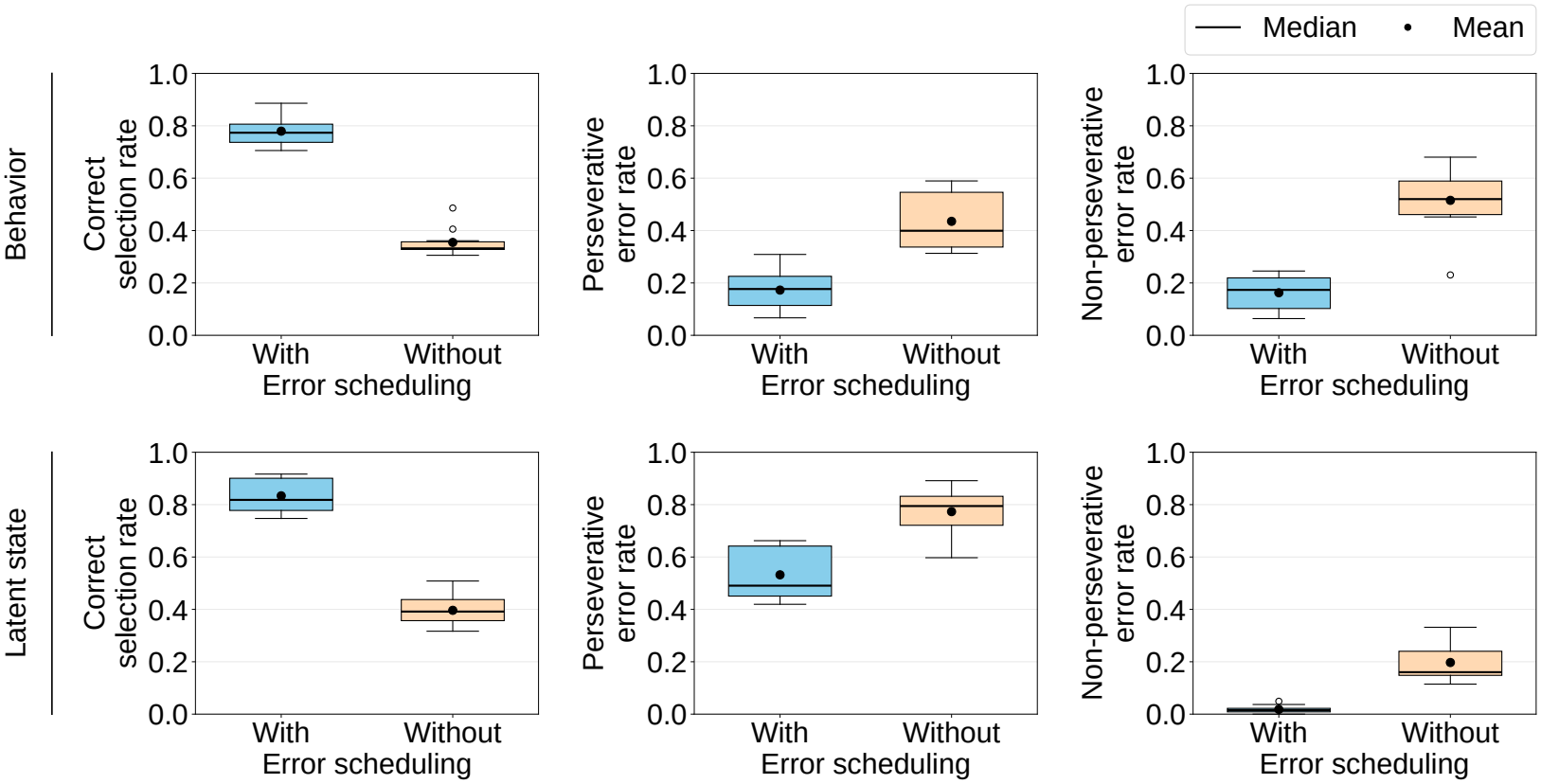
Comparison of evaluation metrics (correct selection rate, perseverative error rate, and non-perseverative error rate) at both the behavioral and latent state levels, between the networks with and without the error scheduling. Each box summarizes 10 networks.

The error scheduling was therefore a condition for the keeping and the switching.

## 4. DISCUSSION

The purpose of this paper is to propose a mechanism behind the balance between stability and flexibility in the WCST through a computational model. Prior models start with a variable for each category, and they treat correct feedback as desirable from the start. We gave the model neither of these, and we modeled the WCST with a hierarchical RNN based on the FEP.

The experiments gave three main results. First, the learning experiments showed that the second-layer latent states formed three clusters in seven of the 10 networks (Figures 7 and 8). Each cluster may correspond to one category. We gave the model neither a category label nor a discrete state for a category, and we removed the auxiliary signals in the testing phase. Second, the test experiments showed that the model kept the estimate after correct feedback and switched it after incorrect feedback in a representative sequence (Figure 9). The posterior mean of the second-layer latent states moved after incorrect feedback. Third, removing the goal-directed process degraded all six metrics (Figures 10 and 11, Table 2). All differences were significant. The keeping and the switching also did not appear without the error scheduling (Figure 12, Table 3). They appeared only when we added the error signals step by step.

The model learned to distinguish the categories, and it selected the card as goal-directed behavior, although we gave it neither a variable for each category nor a preference for correct feedback. Both processes and the balance between stability and flexibility emerged from prediction error minimization based on the FEP. This implies that cognitive processes and this balance in the WCST emerge from the FEP.

### 4.1. FEP-based studies of the WCST

Three earlier studies have applied the FEP to the WCST, and none of them explains the two processes. First, the model of Doulfoukar et al. (2026) holds the categories as discrete states, and it treats correct feedback as desirable from the start. Second, Barceló (2021) interprets the task under prediction error minimization. On this basis, he reviews the EEG and MEG responses to the feedback cues and to the card presentation. This account covers the cortical dynamics, but it implements no computational model. Third, Goto et al. (2025a) formulate the card selection in the same way as the present study. Their model takes correct feedback at future time steps as the target signal, and it generates the card selection by minimizing the error between this target and the predicted feedback. That model, however, simplifies the task in three ways. It sees one card instead of five. It selects one of the card’s feature values instead of one of four target cards. It receives the correct category as an input signal during learning.

The proposed model removed all three simplifications. It used the five cards of the standard WCST, and it selected one of the four target cards. It never received the correct category itself. During learning, it received the card features of the correct category, but these features were not the category itself. We removed them in the testing phase. The present study is therefore the first to show, within the FEP, that the two cognitive processes and the balance between stability and flexibility emerge from prediction error minimization in the standard WCST.

### 4.2. Computational studies of the WCST

We next place the present study among computational models of the WCST outside the FEP. The first group assigns one unit to each category. Levine and Prueitt (1989) give one bias unit to each of color, number, and shape. These units gate the card features, and correct or incorrect feed-back changes their activity. Dehaene and Changeux (1991) give one unit to each category. Incorrect feedback weakens the synapses of the currently active unit, and activity then moves to the unit of another category. Berdia and Metz (1998) keep this design and reproduce the WCST performance of patients with schizophrenia, and Kaplan et al. (2006) reproduce both perseveration and distractibility. Monchi et al. (2000), Caso and Cooper (2017, 2020), and Kajić and Stewart (2021) map the same design onto the structure of neural circuits. The second group assigns a continuous quantity to each category and updates this quantity from the feedback on each trial. Bishara et al. (2010) update the attention paid to each category, and Steinke et al. (2018) apply this model to Parkinson’s disease. Steinke et al. (2019) then add levels of control, and Steinke et al. (2020) run model-based and model-free reinforcement learning in parallel. Steinke and Kopp (2020) review this line of work. D’Alessandro et al. (2020) update the posterior probability of each category by Bayes’ rule instead. The third group adds inner speech and metacognition to the same design. Granato et al. (2020) let an agent manipulate its own representations with inner speech, and Granato and Baldassarre (2021) formalize this manipulation. Granato et al. (2022) then apply the model to autism, Granato and Baldassarre (2024) extend the theory toward consciousness, and Granato et al. (2026) model the metacognitive version of the task.

All of these models start with two things. First, they have a variable for each category. The variable appears as a unit, as a continuous quantity, or as a posterior probability. Second, they treat correct feedback as desirable. They strengthen or keep the current category after correct feedback, and they move to another category after incorrect feedback. They never learn either the category variables or the preference for correct feedback. The model of Liu and Wang (2024) differs in that it acquires the category representation through learning. However, the learning targets are always the correct category and the correct card selection, and this model also does not generate the card selection as goal-directed behavior.

The proposed model differed on both points. First, the three categories appeared as three regions of a continuous latent space, and prediction error minimization formed these regions. Second, the same minimization generated the card selection as goal-directed behavior. The two cognitive processes and the balance between stability and flexibility emerged from one principle.

### 4.3. Computational studies of cognitive flexibility

We now turn to computational studies of the balance between stability and flexibility. Within the FEP, researchers have studied this balance as the arbitration between goal-directed and habitual behavior (Pezzulo et al., 2015, 2018; Parr et al., 2023). In particular, Proietti et al. (2025) propose a hierarchical active inference model in which a meta-cognitive level sits above the behavioral level. The meta-cognitive level controls a parameter that weights goal-directed control against habitual control. It selects its policies by minimizing expected free energy over its own generative model, which is separate from the model of the behavioral level.

Outside the FEP, Rougier et al. (2005) train a neural network model with a prefrontal-cortex-like component whose content is updated by a gating mechanism. Through training on several tasks, this component develops abstract, rule-like representations. They call these representations rules without symbols. Maniadakis and Tani (2009) and Maniadakis et al. (2012) use a genetic algorithm to evolve a continuous-time recurrent neural network (CTRNN) on tasks similar to the WCST. Their networks encode each rule as a neurodynamic attractor and switch rules by moving between attractors. However, their fitness function measures how well the network switches. Correct switching is therefore built into the learning objective.

Another line of computational research concerns the course of learning. Limits on resources or inputs during learning can help a system build complex structure. Elman (1993) trains an RNN on a complex grammar. The network fails to learn when it receives the full grammar from the start. It succeeds only when the input or the processing capacity grows step by step. Elman argues that such limits are not an obstacle but a condition for learning complex domains. Machine learning uses the same idea under the name of curriculum learning (Bengio et al., 2009). Here the examples run from simple to complex, and this order improves generalization. Our error scheduling belongs to this family. It added the error dimensions step by step, and it thereby controlled how complex the learning task was at each stage.

The present study has three characteristics in relation to these lines of work. First, unlike Proietti et al. (2025), our model needed no meta-cognitive level that controls behavioral parameters. Every layer minimized the same free energy, and the keeping and the switching emerged from the goal-directed process. Second, unlike Maniadakis and Tani (2009) and Maniadakis et al. (2012), our learning objective did not include category switching or switching performance. The switching arose from prediction error minimization based on the FEP instead. Third, in line with Elman (1993) and Bengio et al. (2009), the keeping and the switching did not occur unconditionally. They depended on the step-by-step addition of the error signals. This suggests that developmental constraints matter for acquiring cognitive flexibility.

### 4.4. Neuroscientific studies of cognitive flexibility

We now consider the results in light of neuroscientific evidence. The first result is that the model learned to distinguish the categories in the second-layer latent states. Wallis et al. (2001) show that single neurons in the monkey prefrontal cortex code abstract rules rather than specific stimuli. Mansouri et al. (2006) use a task closer to the WCST. In their task, no cue presents the correct rule, and the reward and error signals do not point to the rule either. Rule-related activity still arises in the monkey dorsolateral prefrontal cortex. Neural activity therefore supports internal rule representations during WCST-like tasks. Similarly, Freedman et al. (2001) show that prefrontal neurons come to hold category-related representations through training. Our results suggest that prediction error minimization based on the FEP can explain how such separated representations arise.

The second result is that the model kept the estimate after correct feedback and switched it after incorrect feedback. In Figure 9, the posterior mean of the second-layer latent states shifted for a short time after incorrect feedback. Nakahara et al. (2002) compare humans and monkeys with event-related fMRI on a modified WCST. In both species, activity in the posterior ventrolateral prefrontal cortex rises for a short time at the moment of category switching. Also, Monchi et al. (2001) separate the stages of the WCST with event-related fMRI. The ventrolateral prefrontal cortex, the caudate nucleus, and the mediodorsal thalamus become active only when a participant receives negative feedback and has to switch. Our result may relate to these reports. Also, the model adjusted its current card selection by minimizing the error between correct feedback and the feedback it predicted at future time steps. Holroyd and Coles (2002) report a related pattern in humans. In their learning task, participants have to work out the correct response to each stimulus from reward and penalty feedback. When the correct response is learnable, the error-related negativity (ERN) is larger at feedback than at the response early in learning, and this relationship reverses as learning goes on. A prediction about the outcome is therefore already present when the participant responds.

The third result is that appropriate switching required the step-by-step addition of error signals. The networks performed better with the error scheduling (Figure 12). Human development shows a similar course in both behavior and neural activity. For example, Moriguchi and Hiraki (2009) use the dimensional change card sort (DCCS) task, in which children first sort cards by one rule, such as color, and then have to switch to another rule. Three-year-olds often fail this switch, and five-year-olds perform it almost perfectly, as adults do. Their near-infrared spectroscopy (NIRS) data also show a difference in the right inferior prefrontal region. Among three-year-olds, most children who switch successfully keep this region active throughout the task, and none of those who perseverate do. A similar developmental course appears in the WCST. Chelune and Baer (1986) test 105 children aged 6 to 12 years. Perseverative errors decrease with age, and performance reaches the adult level at about 10 years. These results suggest that appropriate switching in the WCST depends on a course of learning rather than on a mechanism present from the start. Elman (1993) and Vogelsang et al. (2024) argue that limits early in development act as scaffolding rather than as obstacles. Our results support this view. Two differences from human development remain, however. First, age changes many things at once, and our schedule changed only the order of the error signals. Second, no one chooses that order for a child, and we chose it for our model.

### 4.5. Implications for machine learning

Large language models (LLMs) have improved rapidly, and researchers now evaluate their cognitive abilities with psychological tests designed for humans. Researchers call this field machine psychology, and it has reached the WCST. Li et al. (2024) include the WCST in a benchmark of seven tasks and measure how a model revises its belief after an unexpected outcome. Hao et al. (2026) run visual LLMs on the task. With chain-of-thought prompting and textual input, the models achieve or surpass human-level set-shifting performance. Our earlier work also analyzes the card selection of LLMs on the task (Goto et al., 2025b). These studies either measure the performance or infer the mechanism from the outputs. The present study instead describes the card selection in the WCST in terms of a computational principle, pre-diction error minimization. This description may serve as common ground for comparing card selection mechanisms across classes of models, including LLMs.

### 4.6. Limitations and future work

This study has two main limitations. First, we tested the model only in simulation. We did not compare its performance quantitatively with human performance on the WCST. Second, we do not yet know which brain regions correspond to each part of the model.

Future work should address these two limitations. It should also extend the approach to other tasks. Task-switching tasks and working-memory tasks also probe the stability–flexibility dilemma. We aim to test whether an FEP-guided developmental process can explain how people respond in those tasks as well.

